# Characterization of habitat requirements of European fishing spiders

**DOI:** 10.1101/2020.12.13.422580

**Authors:** Lisa Dickel, Jérémy Monsimet, Denis Lafage, Olivier Devineau

**Affiliations:** Department of Forestry and Wildlife management, Inland Norway University of Applied Sciences, Campus Evenstad, Koppang, Norway; Centre for Biodiversity Dynamics, Department of Biology, Norwegian University of Sciences and Technology (NTNU); UMR CNRS 6553 ECOBIO, Université de Rennes, Rennes, France; Department of Environmental and Life Sciences/Biology, Karlstad University, Karlstad, Sweden

**Keywords:** *Dolomedes*, Pisauridae, detectability, red listed species, conservation

## Abstract

Wetlands are among the most threatened habitats in the world, and so are their species, which suffer habitat loss due to climate and land use changes. Freshwater species and arthropods receive little attention in research and conservation, and the goals to stop and reverse the destruction of wetlands published 25 years ago in a manifesto by the Union of Concerned Scientists have not been reached. In this study, we investigated the occurrence and habitat requirements at two spatial scales of two species of European fishing spiders *Dolomedes*, which rely heavily on declining wetland habitats in Sweden and southern Norway.

We collected occurrence data for *Dolomedes plantarius* and *Dolomedes fimbriatus*, using a live-determination-method. We modelled the placement of nursery webs to describe fine scaled habitat requirements related to vegetation and microclimate. Using a machine learning approach, we described the habitat features for each species, and for co-occurrence sites, to provide insight into variables relevant for the detectability of *Dolomedes*.

We found that habitat requirements were narrower for *D. plantarius* compared to *D. fimbriatus*; that the detection of nursery webs can be affected by weather conditions and that nursery placement is mostly dependent on the proximity to water, the presence of *Carex sp.* (Sedges) and of crossing vegetation structures, and on humidity. Furthermore, co-occurring sites were more similar to *D. plantarius* sites than to *D. fimbriatus* sites, whereby surrounding forest, water type and velocity, elevation and latitude were of importance for explaining which species of *Dolomedes* was present.

We provide a detailed field protocol for *Dolomedes* studies, including a novel live-determination method, and recommendations for future field protocols.

## Introduction

Biodiversity is threatened by anthropogenic land use and climate changes (Sala 2000). Wetlands are among the most threatened habitats, yet they are particularly important ecosystems in terms of climate change mitigation, biodiversity by providing breeding and feeding grounds for many species, and hydrology through flood regulation, water holding bodies, and nutrient retention. Thereby, wetlands are crucial to human existence (De Groot et al. 2006). 87% of wetlands have been lost since the beginning of the 18th century, mainly because of expansion of agriculture and urbanization (Davidson 2014). De Groot et al. (2006), and Hu et al. (2017) estimated that 33% of wetlands were lost due to human activities. The Ramsar Convention (Ramsar Convention Secretariat 2013) and the first world’s scientists warning to humanity (Kendall 1992) formulated wetland conservation as a global goal. However, in a recently published second warning to humanity, the authors stated that not only wetland protection and restoration goals were reached, but that wetlands destruction and loss have proceeded (Finlayson et al. 2019) Conservation priorities are mostly determined through variable and dynamic human value (Lindenmayer and Hunter 2010), which has led to unequal conservation efforts across habitats and taxa, with groups like invertebrates (Clark and May 2002, Finlayson et al. 2019) and freshwater/wetland species being particularly neglected (Darwall et al. 2011). Other issues interact with and add on this: the difficult accessibility of wetlands and the low detection of invertebrates (Noreika et al. 2015) cause bias across habitats and taxa. Yet, according to a review by Kellner and Swihart (2014), few studies accounted for imperfect detection, and even less often in invertebrate studies than in studies of other taxa (Kellner and Swihart 2014)..

Although Clark and May (2002) recognized the taxonomic imbalance of research, basic knowledge is still missing to inform conservation of wetlands’ invertebrates. This knowledge is lacking for the two European fishing spiders of the *Dolomedes* genus, namely the raft spider *Dolomedes fimbriatus* and the great raft spider *Dolomedes plantarius*. Both species are semi-aquatic, forage on land as well as on water, and build their nursery webs close to or even above the water surface (Gorb and Barth 1994, Duffey 2012). The detection of both species is difficult due to their lifestyle, which includes fleeing behavior on and under the water surface when disturbed (Gorb and Barth 1994). *Dolomedes* do not build prey-webs, which makes it even more difficult to detect. Like other members of the family Pisauridae (Stratton et al. 2004), *Dolomedes* build nursery webs, which are a convenient sign of presence during the reproductive season, thus facilitating their detection. Females are found close to their nursery webs, which is useful for identification, mainly because only adults can be identified with certainty by inspecting their genitals. Further, the placement of nursery webs functions as an important indicator for *Dolomedes* habitat determining reproductive success and survival. habitat determining reproductive success and survival.

Habitats of both species are declining because of anthropic transformation and draining of wetlands (van Helsdingen 1993, Hu et al. 2017, Finlayson et al. 2019). While *D. fimbriatus* is relatively common (Duffey 2012), *D. plantarius* is much rarer, and is one of the few red-listed spiders in Europe, despite its fairly broad distribution range (Leroy et al. 2013). Naturalist observations suggest that *D. plantarius* has more specific habitat requirements than *D. fimbriatus* (Duffey 2012). Habitat loss for might have more severe consequences for *D. plantarius,* which has less potential to adapt, thus making it a species of conservation interest (Smith 2000). Investigating the population decline is difficult, as historical distribution data of *Dolomedes* are scarce (Duffey 2012). Some authors suggest that there may be denser populations of *D. plantarius* than known, especially in the less monitored areas in eastern Europe (Ivanov et al. 2017). Additionally, misidentifications of the two species were common in the first half of the 20th century, when body color was used for determination, although it is not a reliable indicator for the discrimination of both species (Bonnet 1930, van Helsdingen 1993). Little monitoring combined with potential misidentifications and difficult detection of *Dolomedes* caused an overall lack of knowledge about the distribution and status of the species. Recent observations indicate that co-occurrence, which was considered rare or even impossible, might be more frequent than previously thought (Ivanov et al. 2017).

In this study, we contribute to further characterizing the habitat requirements of the two European *Dolomedes* species. Based on naturalist observations by van Helsdingen (1993) and (1995, 2012), we expect *D. fimbriatus* to be more flexible than *D. plantarius* in its habitat requirements regarding the presence of water, and the specific characteristics of the aquatic habitat. We also expect *D. fimbriatus* to occur at higher altitudes and latitudes due to less restricted temperature requirements, and to tolerate more acidic terrestrial habitats, which would facilitate its presence in mires, bogs, and near coniferous forest. coniferous forest.

## Materials and Methods

### Study Area and Site Choice

In order to find potential *Dolomedes* habitats, we chose our study sites based on prior observations extracted from the Global Biodiversity Information Facility (GBIF: The Global Biodiversity Information Facility 2019) using the R package rgbif (Chamberlain and Boettiger 2017) and based on the habitat suitability map of *D. plantarius* from Leroy et al. (2014). Because the resolution of the suitability map and the accuracy of the GBIF positions were too low for our purpose, we selected sampling areas within the highly suitable habitat and close to the GBIF positions based on information from the literature.

We chose water bodies with riparian vegetation and other types of wetlands (bogs, fens, meadows) for data collection (van Helsdingen 1993, Duffey 1995, 2012). Because the model by Leroy et al. (2014) is only valid for *D. plantarius*, we assessed the potential suitability for *D. fimbriatus* based on the visual impression we had of the wetland, during a visit. The selected locations and the detected species are shown in Figure 1.

**Fig. 1:**
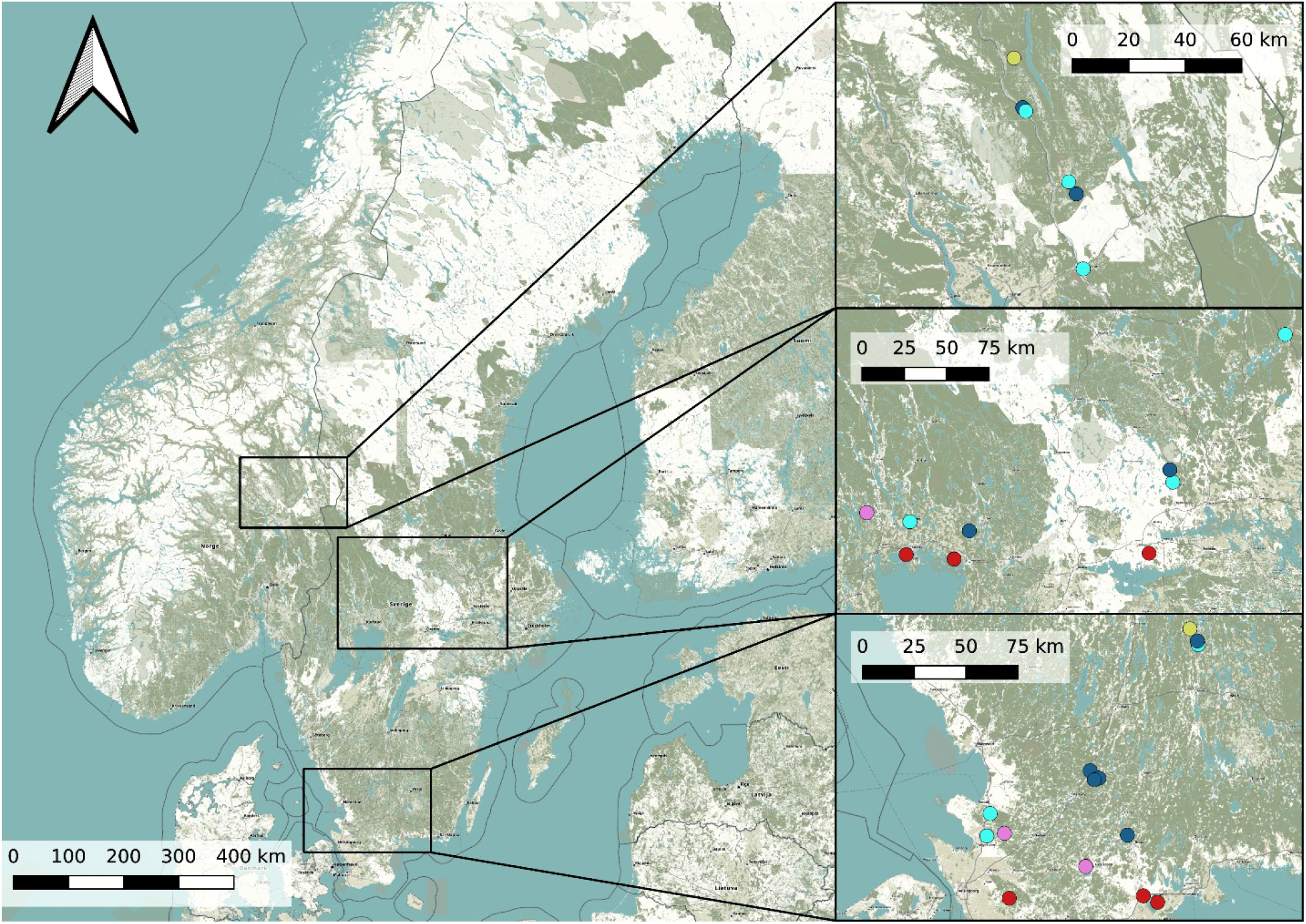
Overview map of the study area in Sweden and Norway (left) with detail maps of the three study areas (right). Dots represents the study sites (Purple: both species; red: *D. plantarius*; Light blue: *D. fimbriatus*; blue: absence sites; beige: *Dolomedes sp.*) The background map is obtained from OpenStreetMap.

### Data Collection

We collected and geo-referenced all data using the data collection software KoBoToolbox (KoBoToolbox 2020). We collected habitat data at site and microhabitat scales. We determined and recorded species of *Dolomedes* at the site scale. We delimited each study site by its natural borders, or, if too big, after five transects; i.e. 40 m along the water body, see transect description below. after five transects; i.e. 40 m along the water body, see transect description below.

#### Site Scale Data

Since the detectability of free-ranging spiders varies with weather conditions (Noreika et al. 2015), we recorded temperature and wind speed, and visually classified rain and clouds at the beginning of each field work session. In case of wind (Beaufort scale > 3, equivalent to 12-19 km/h wind speed) or rain, we did not attempt to detect the spiders, to keep detection conditions equal.

We searched for nursery webs and spiders for 20 min on each site. If possible, we searched the edge of the vegetation both visually and by sweep-netting, while wading through the water. If entering the water was not possible (e.g., due to quality of the substrate, the strength of stream or water depth), we moved carefully across the riparian vegetation to the contact zone of marginal vegetation and water and applied the same search strategy. We found most adult females in nursery webs or in the nearby vegetation, or on the water. We captured the spiders in a glass container. If the spider dived, we caught it with a fishing net (mesh size approximately 0.9*0.3 mm) from the water and transferred it into a glass container.

Once inside the container, we determined the species by pressing the individual gently with a soft sponge against the glass, to inspect the epigyne or pedipalps (A picture of the identification process is available in Appendix A). We released all spiders after identification. If we detected only nursery webs but no spider on a site, we discarded the data, as the nursery web of *Pisaurida mirabilis* cannot safely be distinguished from the web of *Dolomedes*.

We collected variables regarding vegetation type, land use and surroundings at site level (Table 2). As *Dolomedes* are semi-aquatic species, measurements on plot level were concentrated around the water body or in the ‘wet center’ of study sites without open water, from where we drew transects for further data collection plots.

#### Microhabitat Data

Within each site, we placed up to five transects to systematically arrange sampling plots (Fig. 2) to collect microhabitat data. If open water was present, we placed the transects perpendicular to the water body and 10 m apart. If no open water was present, we placed transects along a wet to dry ground gradient. If no gradient was detectable, the transects started (at random) from a habitat edge, to represent the site of interest. We recorded the applied sampling type for each site.

**Fig. 2:**
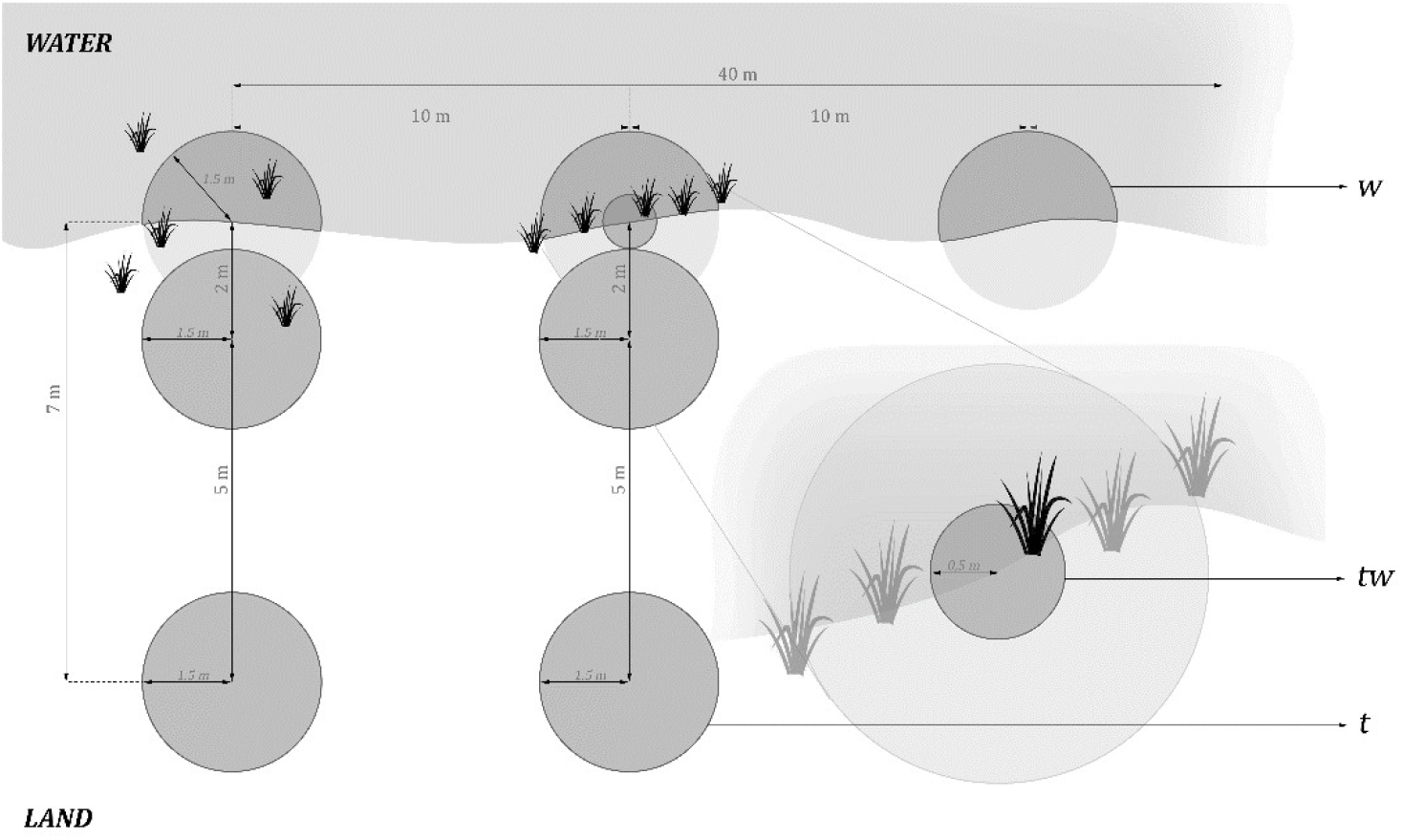
Arrangement of plots transects in a site with open water, i.e. with aquatic plots, a higher density of plots close to the water, and with an additional small terrestrial plot for the narrow riparian vegetation area (middle).

Along each transect, we collected plot-scale data for one aquatic plot (if water was present) at the beginning of each transect and terrestrial circular plots (radius = 1.5 m). Terrestrial plots were located at two, seven and twelve meters from the water edge based on test sites to represent the gradient from aquatic to terrestrial habitat. The focus on the shore-area (or the wettest area in the site) is reflected by the higher density of plots close to the water (Figure 2). We collected data in one to three terrestrial plots, depending on the width of the site. When the riparian vegetation was limited to a few centimeters by the water edge, we included a fourth half-circle (r = 0.5m) terrestrial plot with its center at the water edge to represent the vegetation (see tw plot in Figure 2). The shape of the additional plot differed from the others to avoid plots overlapping. We collected percent cover data for the five most relevant plant species according to literature (*Carex spp*., *Juncus spp*., *Typha spp*., *Phragmites spp.* and *Sphagnum spp.*) on the Braun-Blanquet scale (Westhoff and Van Der Maarel 1978), which we later reduced for modeling purposes (Table 3). Furthermore, we collected structural and microclimate variables (Table 2). We collected the same measurements around the nursery webs, which we searched for in the entire site. We extracted site elevation after data collection from a digital elevation model (EEA 2018).

### Statistical Analysis

We prepared and analysed all data in R (R Core Team 2020), and R Studio (RStudio Team 2012). We followed the protocol for data exploration by Zuur et al. (2010) and used the tidyverse framework for data exploration and preparation (Wickham et al. 2019). We standardized all continuous variables to facilitate model convergence and interpretation.

#### Site Scale Analysis

In order to investigate differences among habitats occupied by the two the occupied habitats of both species, we compared sites in which only *D. fimbriatus*, only *D. plantarius*, both species or no *Dolomedes and none of both species* were detected by using flexible discriminant analysis (FDA) (Hastie et al. 1994) using the R package ‘mda’ (Hastie et al. 2013) We used the *Dolomedes* species detection; i.e. *D. fimbriatus, D. plantarius,* both species, or no *Dolomedes* detected, as the response variable in this supervised machine-learning-model. We considered surrounding landscape and forest type, latitude, elevation, water type, water speed, water clearness, and vegetation type as predictors (Table 1).

**Table 1:** Braun Blanquet scale and simplification used in this study.

| Percentage | Number of occurrences | Original Braun-Blanquet category | Simplified category |
| --- | --- | --- | --- |
| 0 | 0 | no | 0 |
| <1 | 1 | r | 0 |
| <1 | 43953 | + | 0 |
| <5 | 6-50 | 1 | 1 |
| <5 | >50 | 2m | 1 |
| 5-15 |  | 2a | 1 |
| 26-25 |  | 2b | 1 |
| 26-50 |  | 3 | 2 |
| 51-75 |  | 4 | 3 |
| 76-100 |  | 5 | 4 |

**Table 2:** Levels and explanations of variables on landscape/ site scale

| Variable name | Levels |
| --- | --- |
| Surrounding | Infrastructure/forest/other |
| Surrounding forest | Deciduous/coniferous/mixed |
| Water type | River/bog/lake/creek |
| Water speed | Standing/slow/fast |
| Water clearness | Clear/brown/murky |
| Vegetation type (at site) | Open wet/open dry/deciduous forest/coniferous forest |
| Latitude | Latitude (continuous), DEM |
| Elevation | Elevation (continuous), DEM |
| Clouds (detectability) | Yes/no/partly |
| Wind (detectability) | Measured with anemometer on Beaufort scale |
| Reason visit | Suitable habitat/GBIF/other |

**Table 3:** Levels and explanations of variables on microhabitat/ nursery scale

| Variable | Description/Levels |
| --- | --- |
| *Dolomedes* detection | Spider/nursery web/no |
| Aquatic vegetation | Braun-Blanquet scale |
| Distance to water | No water; 0m; 0.7m; 2m; 7m |
| Humidity | Measured at ground level and 20cm above ground |
| Dominant plant group | Dominant group of plants in plot |
| Horizontal cover | Measured at 10cm; 30cm; 50 cm above ground |
| Maximum height | Maximal height of vegetation in plot, measured with 10cm accuracy |
| Average height | Measured 5 times in random location within plot with 10cm accuracy |
| Tussocks | Tuft of grasses or sedges, measured on Braun-Blanquet scale |
| Large leaves | Yes/no |
| Litter | Yes/no |
| Shade | Yes/no/partly |
| Crossing structures | Abundance of crossing vegetation structures, measured on Braun-Blanquet scale |
| *Carex* spp. | Abundance measured on Braun-Blanquet scale |
| *Juncus* spp. | Abundance measured on Braun-Blanquet scale |
| *Typha* spp. | Abundance measured on Braun-Blanquet scale |
| *Phragmites* spp. | Abundance measured on Braun-Blanquet scale |
| *Sphagnum spp.* | Abundance measured on Braun-Blanquet scale |
| Deciduous plants | Abundance measured on Braun-Blanquet scale |
| Aquatic vegetation | Yes/no |
| Nursery web detected | Yes/no |
| Nursery height | Height of web above ground or water |
| Nursery plant | Host plant of nursery web |
| Number of Nurseries in plot | Number of nursery webs detected in one plot |

In addition, we used a single-season occupancy model (MacKenzie et al. 2002) within the unmarked package (Fiske and Chandler 2011) to predict the nursery detection probability pooled for both species. We used the nursery and plot-scale data as spatial replicates, as an alternative to the usual temporal replicates/detection attempts. We considered weather, microclimatic variables (wind, cloudiness, rain, shade), vegetation structure and sampling related variables as potentially influencing detectability.

#### Microhabitat Characteristics Around Nursery

We modeled nursery presence/absence for sites in which *Dolomedes* presence was verified and in which we found at least one nursery web. Thereby we ensured that the sampling was not temporally unsuitable or the site generally unsuitable, which allowed us to model nursery placement within generally suitable sites.

For variable selection and parameter estimation, we fitted a binomial Generalized Additive Model (GAM) by component-wise boosting, using package mboost (function gamboost, Hothorn et al. 2020). Prior to model fitting, we checked variables correlation and we dropped highly correlated variables accordingly (threshold 0.7, Dorman et al. 2012). We did not consider interactions due to the low sample size. From variables humidity at ground level and humidity at 20 cm above ground level, we kept humidity at ground level. We kept average vegetation height, but we discarded the highly correlated maximum vegetation height. We then fitted a regularized model, following the recommendation in Hofner et al. (2018) using all other predictor variables to identify the most relevant predictors and estimate the model parameters. We validated the model using cross validation and present the final model estimates.

To validate the model, we tested the stability of the selected variables via resampling using the package ‘stabs’ (Hofner and Hothorn 2017). Stability selection provides a reliable way to find an appropriate level of regularization, to keep variables with high selection probabilities. In our model, we used standard choices of tuning parameters, with a cut-off of 0.75 and the number of falsely selected base learners tolerated of 1 (Meinshausen and Bühlmann 2010). A conceptual overview of the analyses performed and the data used is presented in Figure 3 and in Appendix C.

**Fig. 3:**
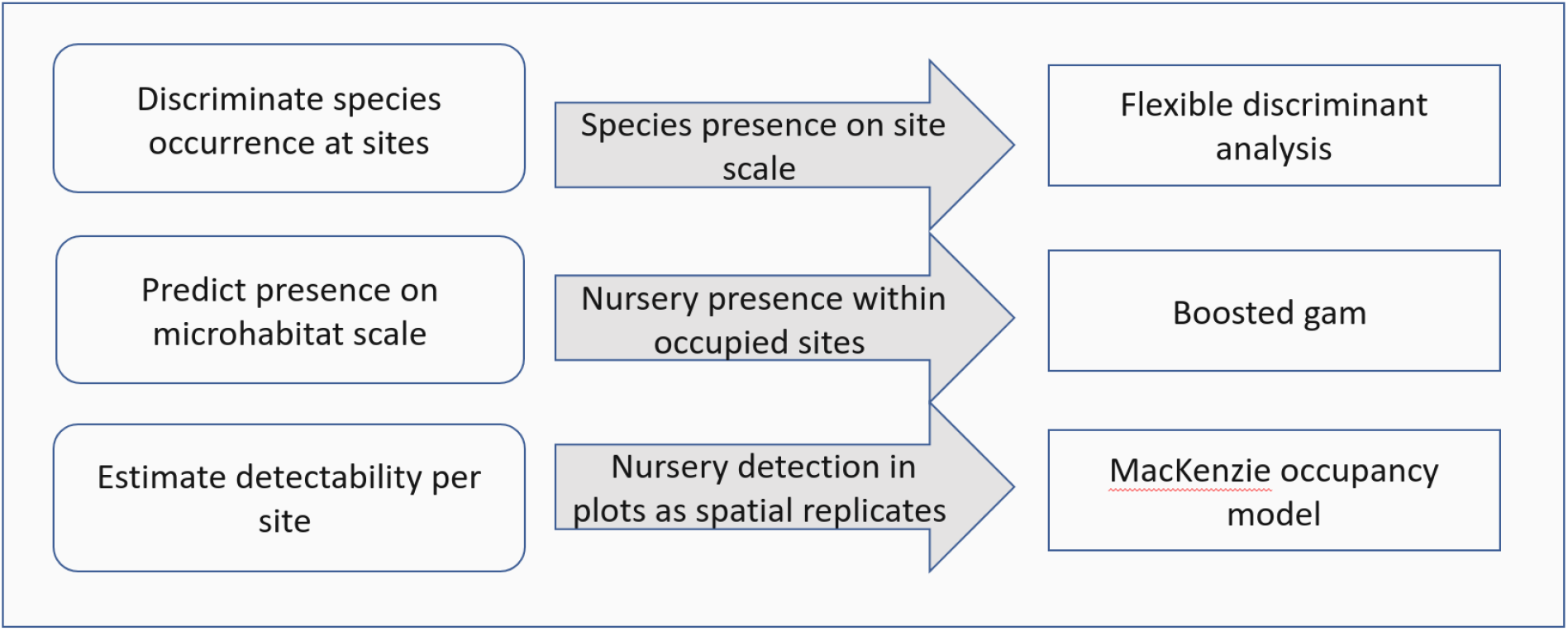
Objective (left), data (middle), and analysis (right) performed in this study and described in the text.

## Results

### Site Scale Habitat Characteristics – Species Specific – Flexible Discriminant Analysis

We detected *D. fimbriatus* alone in 12 sites, *D. plantarius* alone in 6 sites, both species together in 4 sites and none of the two species in 9 of the visited sites (total sites: n= 31, Fig. 1).

The first axis of the FDA explained 82.58%, the second axis 10.34% (Fig. 4). The main variables loading onto the FDA axes, i.e., discriminating best between sites with none/ both / each species, were water type and surrounding forest on the first axis, and water speed on the second axis.

**Fig. 4:**
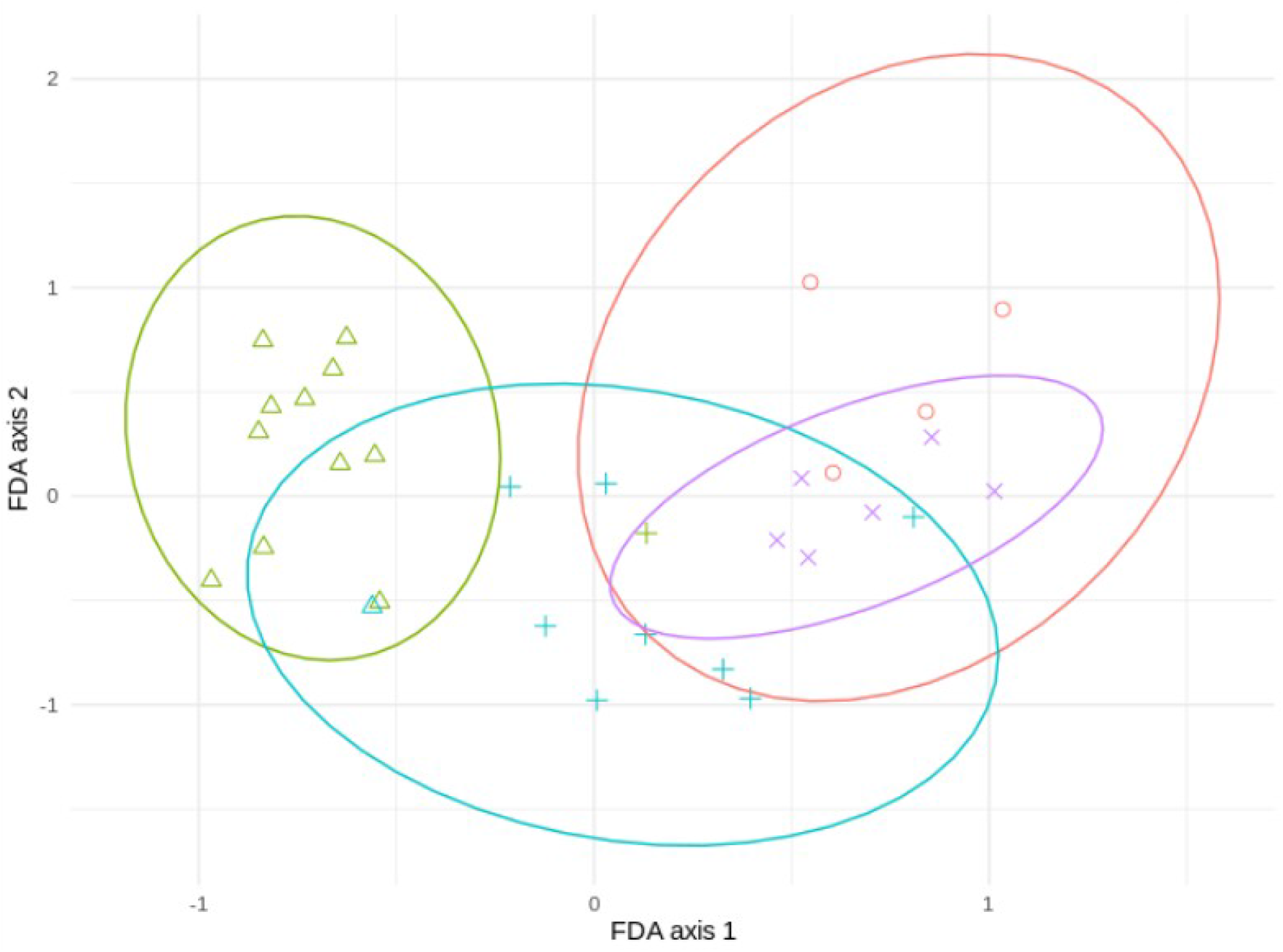
Results of the flexible discriminant analysis (FDA) to separate sites with both species (red), with *D. fimbriatus* only (green), with *D. plantarius* only (purple), or with no *Dolomedes* (blue). Ellipses size indicates uncertainty (95% confidence intervals).

Sites holding either of the two species were well discriminated by the combination of variables loading on the first axis. *D. plantarius* sites were more restricted with respect to the associated habitat variables, and sites with both species overlapped mostly with *D. plantarius* sites (Figure 4).

### Microhabitat Characteristics Around Nursery

In the field, we found a nursery in 35 plots out of 184. The main variables selected in the boosted GAM model (Figure 5) were distance to water (selection frequency = 0.66), *Carex spp.* cover (selection frequency 0.13), crossing structures (selection frequency 0.09), the random effect site ID (0.08), humidity at ground level (selection frequency 0.02) and the intercept (selection frequency 0.01). We found that high abundances of *Carex spp.*, crossing structures, high values of humidity and low distances to water increased the probability of presence of a *Dolomedes* nursery (Figure 6). If water was present, the probability of encountering nursery webs beyond 70 cm away from the water edge was low. There was variation in the probability of finding nursery webs across sites (Figure 6). However, when testing the stability of the selected variables via resampling, only distance to water, the intercept and the random site ID were found to be stable enough, which was most likely caused by the small sample size (Appendix C).

**Fig. 5:**
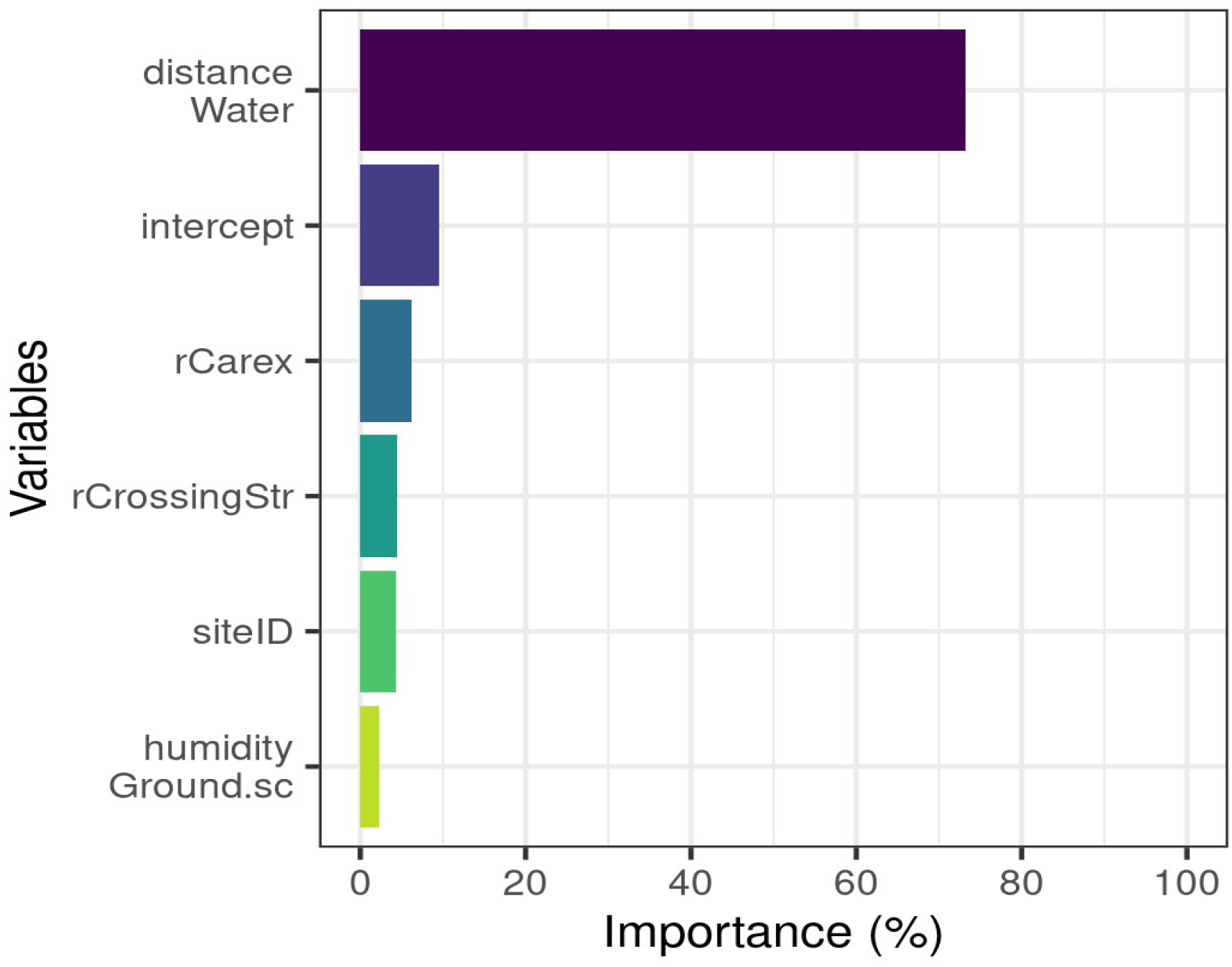
Variables importance for the nursery placement model (boosted GAM). distanceWater: distance to water, rCarex: abundance of *Carex* on simplified Brown-Blanquet scale, rCrossingStr: crossing vegetation structures on simplified Brown-Blanquet-scale, siteID: varying intercept per site ID, humidityGround.sc: humidity at ground level (standardized).

**Fig. 6:**
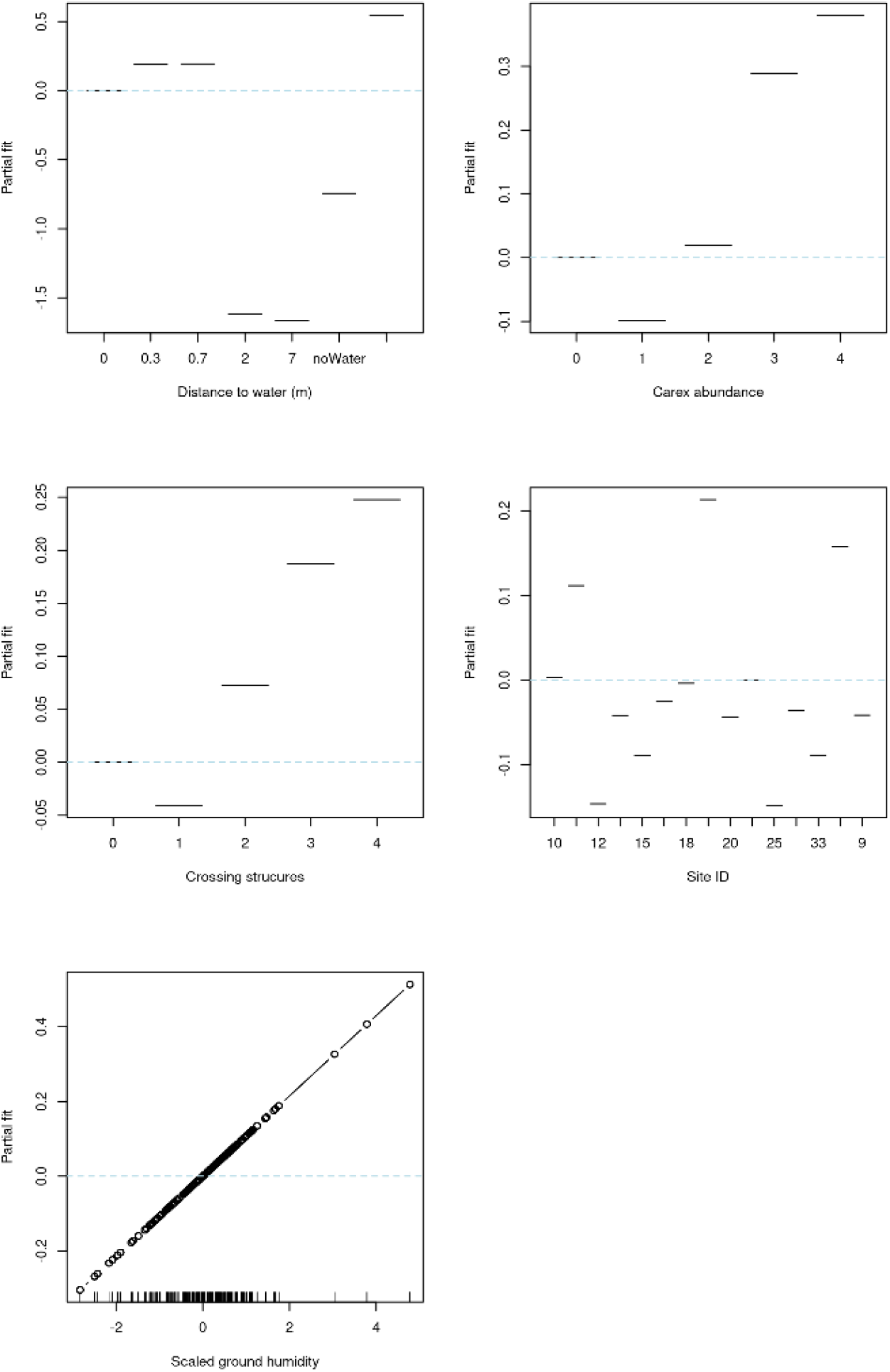
Marginal effects for variables selected in the nursery placement model (boosted GAM).

The detection probability of nursery webs was higher for plots with a high abundance of crossing structures, higher air temperatures, fewer clouds (at the time of data collection), as well as for sites with open water compared to sites without a water body (Figure 4). Model details can be found in Appendix D.

## Discussion

In this study, we found that the two *Dolomedes* present different habitat requirements. We found that *D. fimbriatus* is more generalist than to *D.* p*lantarius.* Forested habitats, and habitats with low pH (as indicated by the presence of species such as *Carex* and *Sphagnum* and the proximity to coniferous forest, Blacklocke (2016)) or without open water appear to still be suitable for *D.fimbriatus*. At the microhabitat scale, nurseries were more likely to be located close to water, and where sedge and crossing structures were present.

The habitat requirements of the two fishing spiders were discriminated mainly by water, represented by the water type and speed, and by the site’s surroundings, both in terms of landscape and forest. We found that *D. fimbriatus* was more tolerant to forested areas, and especially of coniferous forest. Duffey (1995) hypothesized that *D. fimbriatus* can occupy habitat with lower pH values compared to *D. plantarius*. Surrounding coniferous forest, which are dominant in Fennoscandia, may acidify water streams (Blacklocke 2016), thus impacting the pH and potentially restricting *D. plantarius.* We found *D. plantarius* most often at sites with slow-flowing rivers, and we found *D. fimbriatus* most often in bogs. *D. plantarius* was also highly associated with open and slow water, whereas *D. fimbriatus* was less restricted by water conditions. *Dolomedes* can use water as a hunting area and benefit from the use of vibrations at the water surface to detect prey (Bleckmann and Lotz 1987). This close relationship to water, together with the observation of juveniles of *D. fimbriatus* far from the shore, while juveniles of *D. plantarius* are found on the water (Duffey 2012), might result from different hunting abilities between the two fishing spider species.

We found some overlap in, which reflected a spatio-temporal overlap at the site scale. Holec (2000) hypothesized that co-occurrence of both species might only be observed in transitional habitats between sites suitable for *D. plantarius* (ie ponds) and sites suitable for *D. fimbriatus* (ie bogs). This observation is validated for one of the sympatric sites sampled (Figure 1), a fen in the forest. Nonetheless, we hypothesize that the conditions for co-occurrence are less restrictive because similar to (Lecigne 2016), we found two sympatric populations on the vegetation at the shore of a lake (Finjasjön lake, Figure 1). As already hypothesized by van Helsdingen (1993) or Duffey (1995, 2012), our data confirmed the higher degree of association of *D. plantarius* with water compared to D. *fimbriatus*, and in general more substantial restrictions in potential habitats for the former. This suggests that *D. plantarius* is more strictly specialist than *D. fimbriatus*. Even though the separation between generalist and specialist is not clear-cut, species which are ‘rather’ specialist seem to face extinction more often than generalists, especially when facing environmental changes (Colles et al. 2009). The co-occurrence of the two species might be explained by a broader ecological niche of *D. fimbriatus,* which partly overlaps with the niche of *D. plantarius*. The co-occurrence observed hide a possible segregation of both fishing spider species at the microhabitat scale.

At the microhabitat scale, *D. plantarius* might be more dependent on water for its reproductive behavior and nursery placement, which will require further species-specific investigation. Moreover, distance to water and humidity of the ground influenced nursery web placement. This dependency of *D. plantarius* on the water could facilitate cohabitation with *D. plantarius* being spatially segregated towards the shore. Indeed, DeVito et al. (2004) found that spider assemblages of Lycosidae in a microhabitat was influenced by the physiology of the species. We also observed, in two sympatric populations, *D. plantarius* females carrying egg-sac while females of *D. fimbriatus* were already guarding their nursery webs with spiderlings. This might indicate temporal segregation as well, which would also facilitate the co-occurrence of otherwise ecologically close spiders species (Uetz 1977, Fasola and Mogavero 1995). Lastly, the coexistence of wandering spiders at the site scale is usually possible by segregation in micro-habitat use, which is mainly triggered by competition for food (Uetz 1977). An equivalent segregation could occur based on the different diet of the two species, with juveniles of *D. plantarius* being more restricted to water (Duffey 2012).

Within habitats occupied by *Dolomedes*, we found at the microhabitat scale that abundance of sedges (*Carex sp.*) and crossing structures, together with distance to water and humidity, were the most relevant variables for predicting the presence of nursery webs. Indeed, the architecture complexity of the vegetation is important for wandering spiders (Vasconcellos-Neto et al. 2017), and is illustrated here by the positive influence of the presence of crossing structures. We can also hypothesize that spiders benefit from the stiff stems of the sedge more than being taxonomically exclusive to them for placing nurseries. De Omena and Romero (2008) showed that some species which are associated with specific host plants are sometimes mostly dependent on the plant’s architectural structure for hunting and as dwelling. The relation between plant community and plant architecture might then be crucial for fishing spiders, as for predatory arthropods in general (Woodcock et al. 2007). The structural aspect should be for conservation of fishing spiders, e.g. by managing the mowing season.

In this study, our sample was small due to the rarity of the two species, especially in Scandinavia, and due to a narrow temporal window of data collection. At the landscape scale, this small sample, and especially the lack of co-occurrence sites limits the scope of our conclusions about the characteristics of sympatric populations. At the microhabitat scale, repeated visits of the same sites would provide opportunities to refine the occupancy model and to clarify detection issues for these two species. With a better knowledge of nursery timing, other microhabitat studies would also be facilitated. Further data collection at the landscape level would increase knowledge about potential habitat, and investigating water and soil acidity could be helpful to clarify habitat restrictions for *D. plantarius*. Finally, species-specific occupancy modeling could be helpful, as especially *D. plantarius* is likely to dive when disturbed and might be more difficult to detect than *D. fimbriatus*, which might prevent identification of double-species sites.

While the distribution of both *Dolomedes* species might shift towards the North (Leroy et al. 2013, 2014, Monsimet et al. 2020), this shift might be limited by low dispersal abilities and unconnected habitats in Fennoscandia (Monsimet et al. 2020). It is therefore essential to protect both current and future habitats. Conserving both *Dolomedes* species emphasizes the special importance to protect wetlands, in Fennoscandia and elsewhere (Sala 2000, Davidson 2014, Carson et al. 2019). The conservation of the red-listed *D*. *plantarius* might be prioritized as it seems to have narrower habitat requirements than *D. fimbriatus,* which makes it more vulnerable to climate change (Cardoso et al. 2020). Moreover, a reassessment of the outdated IUCN status (World Conservation Monitoring Centre 1996) of the red-listed *D. plantarius* is needed. Management and conservation could here serve and inform each other, as more monitoring of *Dolomedes* could contribute to more protection of wetlands.

To counteract various threats, which spiders currently face, land protection and the management of both land and species, is important (Branco and Cardoso 2020). For efficient management, estimating the local probability of presence of the species is important. Occupancy modeling can help to decide which areas could be necessary to protect and where to apply conservation efforts (McFarland et al. 2012). In this study, the detection probability of nursery webs was higher where abundance of crossing vegetation structures was high and with good weather condition, i.e., optimal temperature and sunny weather. Nonetheless, the use of nursery webs as detection units could be improved by specifying the timing and duration of nursery webs with repeated visits (e.g. weekly) to the same sites and nursery webs (Smith 2000). Monitoring nursery webs also makes it possible to encounter the female spiders, which is especially valuable with the non-invasive sponge-technique we used for identifying the species. Additionally, to estimating the population’s abundance dynamic, the management of population by preserving wet habitat and with abundant crossing structures is essential.

Nature conservation is driven and motivated by a normative component, which is dependent on flexible human values (Lindenmayer and Hunter 2010), causing direct effects on the protection of invertebrates which are traditionally considered less in conservation (Clark and May 2002, Troudet et al. 2017) and might be one of largest obstacles in spider conservation (Branco and Cardoso 2020). For this reason, raising awareness for the conservation of arthropods like fishing spiders is fundamental, e.g. with citizen science projects (Troudet et al. 2017), like SpiderSpotter (“SpiderSpotter” 2020).

## Acknowledgment

We thank Lars Jonsson (Kristianstad University) for his advice and help during the data collection, Andres Ordiz for comments on a previous manuscript version, and Boris Leroy for sharing with us his prediction maps to select sites.

## Data availability

We added new records to Gbif (indirectly, they are on artsobservasjoner and artportalen).

