## Appendix A for "Characterization of habitat requirements of European fishing spiders"

### Appendices

#### *Appendix A: Identification method*

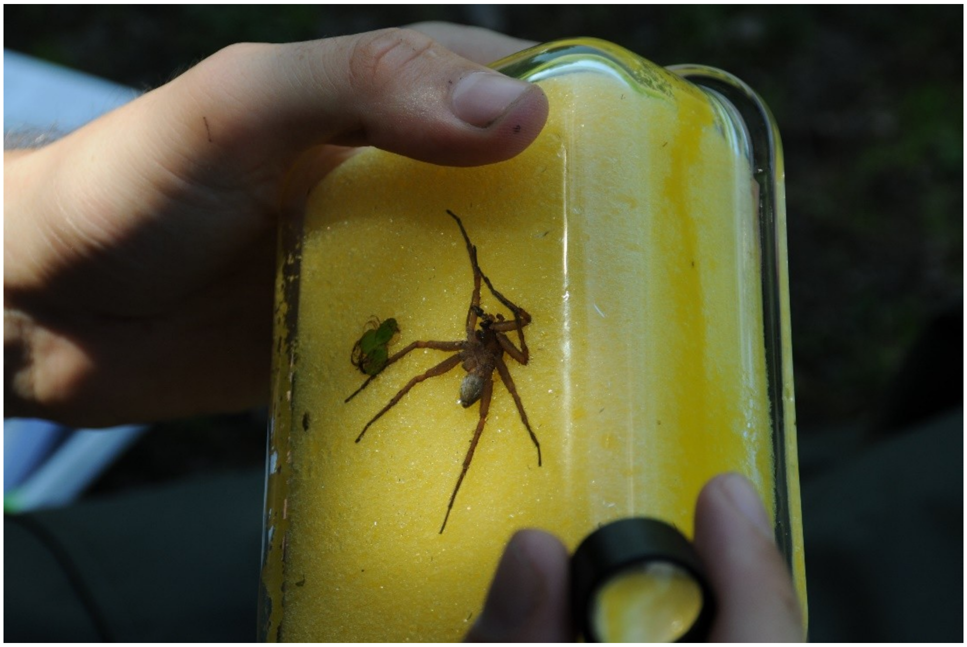

Figure A1: Sponge determination technique for live species identification of *Dolomedes* spiders.

In order to determine the species of *Dolomedes*, we captured the spiders in a glass box, then removed carefully the lid and inserted the sponge (a soft sponge for normal housework activities, prewashed, but dry at the time of use). We then pushed the spider gently to the bottom of the box, paying attention that the spider was in a correct position, with the legs pointing away from the body (important for not harming the spider and for good conditions for determination). We then used a magnifying class to inspect the epigyne or the pedipalps and compared them with illustrations and descriptions of the *Dolomedes* genitals relevant for species identification.
