## Appendix B for "Characterization of habitat requirements of European fishing spiders"

### *Appendix B: Site scale data*

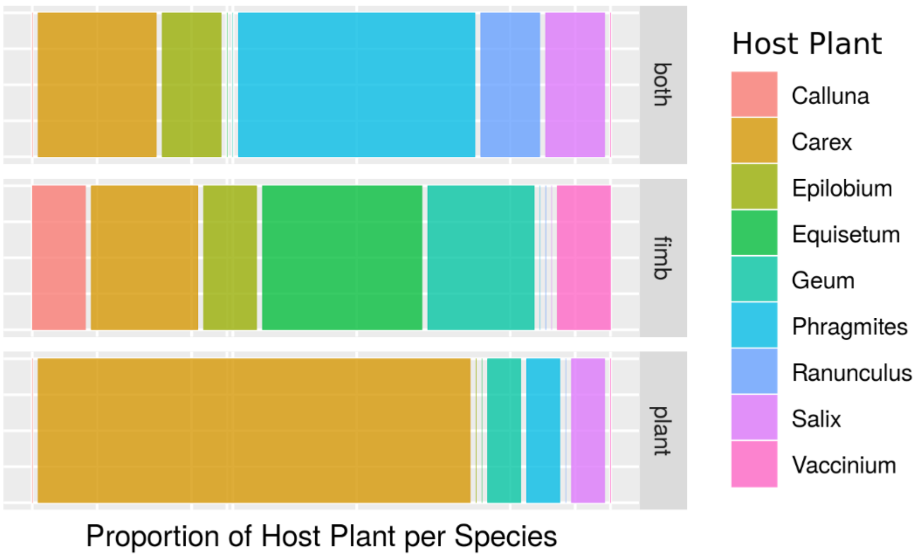


Fig. A1: Frequency of nursery host plants in sites where both species were found (top section: ‘both’), and in sites where only *D. fimbriatus* (middle section : ‘fimb’), or only *D. plantarius* (bottom: ‘plant’) was detected.
