## Appendix C for "Characterization of habitat requirements of European fishing spiders"

### *Appendix C: Nursery placement model (boosted GAM)*

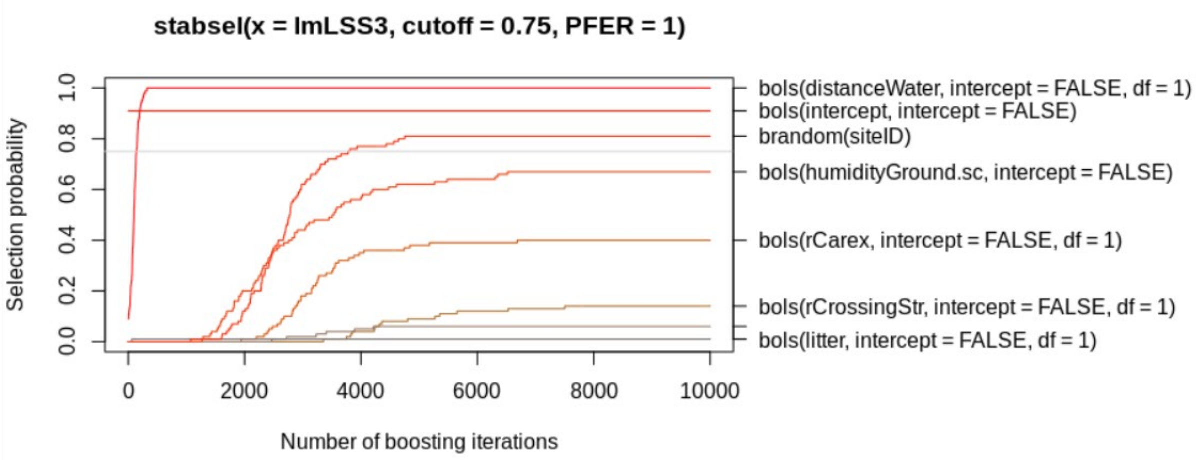


Figure A3: Model validation of boosted GAM for nursery placement, indicating the stability of the selected variables. The steeper the first part of the curve, and the earlier it flattens, the more stable the selection of the variable across resampling, ie the more important the variable. The grey horizontal line indicates the threshold of acceptance.
