## Appendix D for "Characterization of habitat requirements of European fishing spiders"

***Appendix D: Occupancy model***

Table A1: Top five models of the model selection for the occupancy model.

| Model | negLogLike | delta | AIC | AICwt |
|---|---|---|---|---|
| ~ sampling_type ~ 1 | 89.36 | 0.00 | 202.71 | 0.45 |
| ~1 ~ Cattle_grazing + type | 90.95 | 1.18 | 203.9 | 0.25 |
| ~ rCrossingStr + temperature + cloudyness + sampling_type ~ 1 | 92.86 | 3.00 | 205.71 | 0.1 |
| ~ temperature ~ 1 | 89.36 | 4.00 | 206.71 | 0.06 |
| ~ shade ~ 1 | 92.57 | 4.43 | 207.14 | 0.05 |

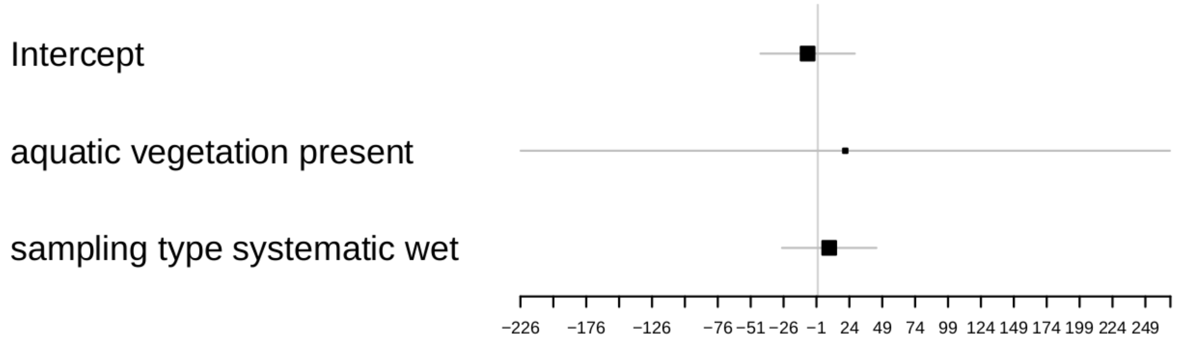


Figure A4: Estimates for occurrence parameters, from the best occupancy model.

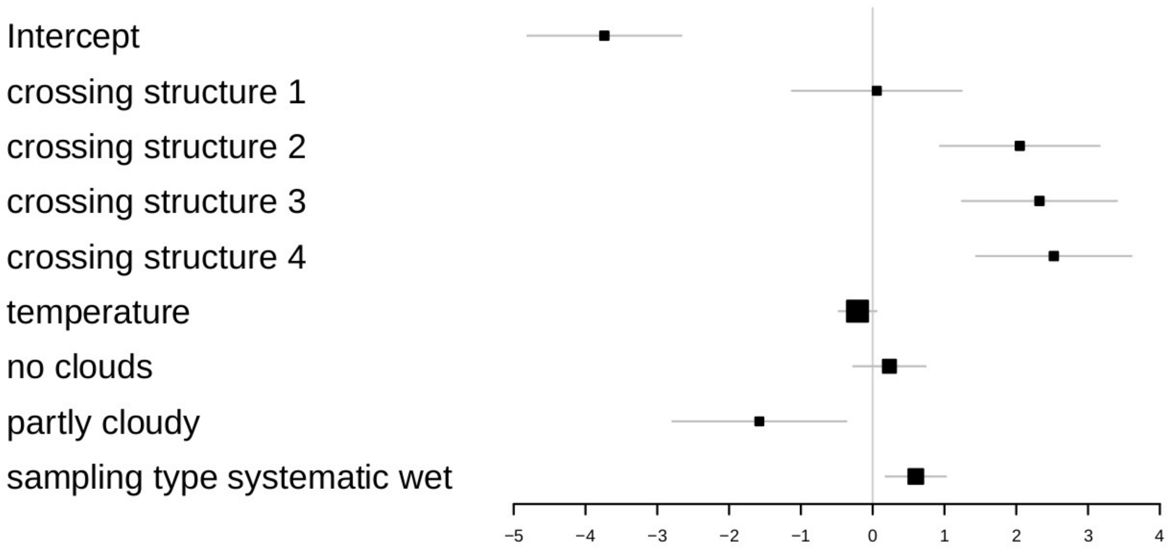


Figure A5: Estimates for detectability parameters, from the best occupancy model.
